# Mathematical modeling to guide experimental design: T cell clustering as a case study

**DOI:** 10.1101/2022.03.30.486434

**Authors:** Harshana Rajakaruna, Vitaly V. Ganusov

## Abstract

Mathematical modeling provides a rigorous way to quantify immunological processes and to discriminate between alternative mechanisms driving specific biological phenomena. It is typical that mathematical models of immunological phenomena are developed by modellers to explain specific sets of experimental data after the data have been collected by experimental collaborators. Whether the available data are sufficient to accurately estimate model parameters or to discriminate between alternative models is not typically investigated. While previously collected data may be sufficient to guide development of alternative models and help estimating model parameters, such data often do not allow to discriminate between alternative models. As a case study we develop a series of power analyses to determine optimal sample sizes that allow for accurate estimation of model parameters and for discrimination between alternative models describing clustering of T cells around Plasmodium liver stages. In our typical experiments, mice are infected intravenously with Plasmodium sporozoites that invade hepatocytes (liver cells), and then activated CD8 T cells are transferred into the infected mice. The number of T cells found in the vicinity of individual infected hepatocytes at different times after T cell transfer is counted using intravital microscopy. We previously developed a series of mathematical models aimed to explain highly variable number of T cells per parasite; one of such model, the density-dependent recruitment (DDR) model, fitted the data from preliminary experiments better than the alternative models, such as the density-independent exit (DIE) model. Here we show that the ability to discriminate between these alternative models depends on the number of parasites imaged in the analysis; analysis of about *n* = 50 parasites at 2, 4, and 8 hours after T cell transfer will allow for over 95% probability to select the correct model. The type of data collected also has an impact; following T cell clustering around individual parasites over time (called as longitudinal (LT) data) allows for a more precise and less biased estimates of the parameters of the DDR model than that generated from a more traditional way of imaging individual parasites in different liver areas/mice (cross-sectional (CS) data). However, LT imaging comes at a cost of a need to keep the mice alive under the microscope for hours which may be ethically unacceptable. We finally show that the number of time points at which the measurements are taken also impacts the precision of estimation of DDR model parameters; in particular, measuring T cell clustering at one time point does not allow accurately estimating all parameters of the DDR model. Using our case study, we propose a general framework on how mathematical modeling can be used to guide experimental designs and power analyses of complex biological processes.

## Introduction

Mathematical modeling is the main, perhaps the only rigorous way to infer and accurately quantify biological processes underlying a specific phenomenon. A typical application of mathematical modeling is to fit a set of alternative mathematical models to an experimental dataset, estimate model parameters, and discriminate between the alternative models [1]. It is fairly common that mathematical models are developed after specific, often preliminary experimental data have been collected; in some cases such an approach precludes the ability to discriminate between alternative models (e.g., [2–5]). In contrast, in randomized clinical trials, it is typical to provide power analyses that drive the sample size of the trial to detect a potential effect of the treatment [6]. Such power analyses are typically the result of a smaller trials that give an estimate of the treatment’s effect size. Power calculations are rare in mathematical modeling-based studies in immunology [7]. By using an example of a specific immunological system here we provide a theoretical framework to perform such power analyses and to determine the sample size for a given type of data that would be required to accurately estimate parameters of a given mathematical model and to discriminate between alternative models.

Several previous studies have shown that vaccinations resulting in generation of a large number of Plasmodium-specific CD8 T cells, are able to provide protection following infection with Plasmodium sporozoites [8–11]. By using intravital imaging, we and others have shown that infection of immune mice with Plasmodium sporozoites results in formation of clusters around sporozoite-infected hepatocytes in murine livers [12–14]; we have shown that formation of such clusters is important in T cell-mediated elimination of the parasites [12, 15]. To study potential mechanisms driving formation of such clusters around infected hepatocytes our previous experiments have been performed in the following manner. First, we infected the mice with a high dose of Plasmodium yoelii sporozoites, expressing green fluorescent protein (GFP). Then 20 hours later we transferred activated fluorescently-labeled Plasmodium-specific CD8 T cells (and in some cases, T cells of irrelevant specificity) to the infected mice, and the number of T cells located within 40 μm of randomly chosen GFP-expressing parasites in different liver areas/mice was determined at 4 hours post-infection [12]. Interestingly, there was a large variability in the number of T cells per parasite; most parasites had no T cells while few parasites had up to 10 T cells [12]. We developed several alternative mathematical models, based on stochastic linear birth-death process modeling framework, predicting the T cell cluster size. The null model, assuming that T cell recruitment into the cluster occurs at a constant rate while exit from the cluster was dependent on the cluster size, did not fit the data well [12]. Two alternative models, density-dependent recruitment (DDR) model (assuming that rate of T cell entry into the cluster depends on the cluster size, **Figure 1**A) and density-independent exit (DIE) model (assuming that per capita exit rate from the cluster declines with cluster size, **Figure 1**B) fitted some of the data with similar quality [12]; however, a subsequent analysis demonstrated that the DDR model was better at describing most of the T cell clustering data including data from experiments of other investigators [16].

**Figure 1:**
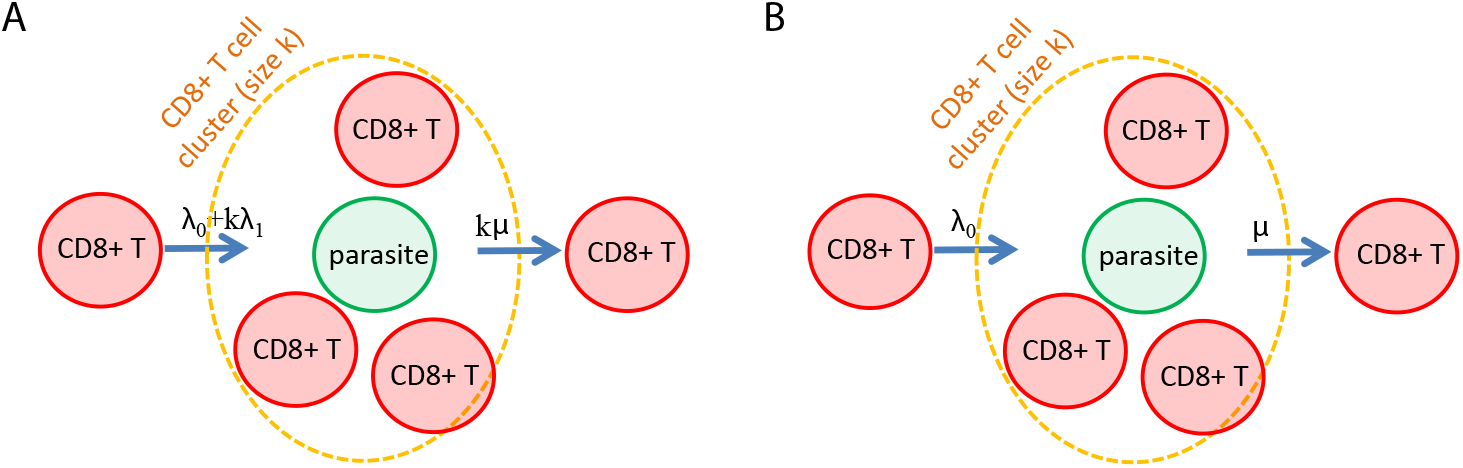
Two competing models to explain variability in the number of Plasmodium-specific CD8 T cells found around individual Plasmodium-infected hepatocytes. In the illustration, the green denotes the parasite and red denotes T cells clustering around the parasite. (A) Density-dependent recruitment (DDR) model, which is similar to the birth-death-immigration model in stochastic processes. (B) Density-independent exit (DIE) model, which is similar to the immigration-emigration model in stochastic processes. Parameters λ_0_, *k*λ_1_, *μ* are T cell immigration, cluster size, *k*-dependent entry, and exit rates, respectively, in eqns. (1)–(3).

In our experiments T cell clustering around the infected hepatocytes was measured in two different ways. (1) In cross-sectional (CS) data, parasites located in different parts of the livers of individual mice were imaged at one or two time points, and the number of T cells per parasite was determined. Such data can be relatively easily generated because such imaging requires taking only snapshots of the volume around an infected hepatocyte that can be performed rapidly in one or several mice for tens to hundred of parasites. In contrast, (2) T cell clustering can be followed longitudinally (LT) around individual infected hepatocytes over time by recording the number of T cells found at different time points after T cell transfer [16]. The LT data are much more difficult to collect because only few parasites can be followed over time in each mouse and for relatively short times (e.g., 2-4 hours). While we could fit the DDR and DIE models to the CS and LT data obtained in our previous pilot experiments [15, 16], several issues remained unexplored: specifically, (1) whether the available data were sufficient to accurately estimate the model parameters, (2) how the number of time points, (3) the number of parasites surveyed, and (4) the methods of data-collection (CS vs. LT data) impact the accuracy of parameter estimates. Here, we used results from our earlier, preliminary experiments and mathematical model-based analyses, performed stochastic simulations of cluster formation using parameter estimates found in analyses of previously published data. For such simulations, we performed several power analyses to determine sampling frequency and number of parasite need to be surveyed experimentally for accurate estimation of model parameters, and the ability to discriminate between alternative (e.g., DDR and DIE) models in the follow up future experiments.

## Materials and methods

### Mathematical models

We propose the following framework of random processes to describe the probability *P_k_* (t) to observe *k* T cells around a given parasite at time *t* after T cell transfer. This method is a follow up on our previous modeling experiments [12], in which the number of T cells found near individual Plasmodium-infected hepatocytes was highly variable.

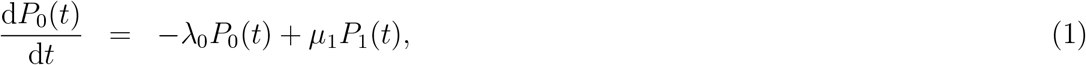

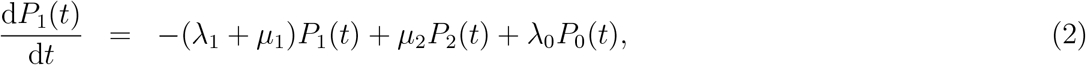

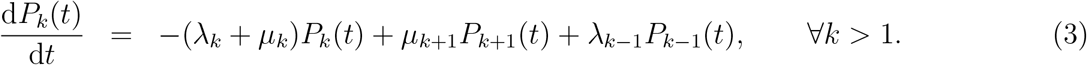

where λ*_k_* and *μ_k_* are the rate at which T cells enter and exit the cluster of size *k*, respectively. In the simplest, random entry-random exit model, entry into the cluster and per capita exit rate from the cluster of size *k* are constants λ_*k*_ = λ_0_ and *μ_k_* = *kμ*, respectively. This model typically did not fit experimental data well with few exceptions, e.g., random clustering of activated T cells that are specific to irrelevant antigens [12, 16].

To explain clustering of T cells in most of other conditions, we proposed two alternative models: (1) The density-dependent recruitment (DDR) model entry into the cluster is amplified by the number of T cells already present in the cluster so λ_*k*_ = λ_0_ + *k*λ_1_ with per capita exit rate being constant, *μ_k_* = *kμ* (**Figure 1**A). (2) An alternative, density-independent exit (DIE) or “retention” model, entry into the cluster is constant but per capita exit rate declines with cluster size *k*, *μ_k_* = *kμ*/*k* = *μ* (**Figure 1**B). In our previous studies, we either fitted the steady state distribution of cluster sizes, 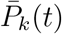, or numerical solutions of the model (eqns. (1)–(3)) to the experimental data [12, 16] to estimate the model parameters. Given our experimental design, the initial condition for all models is *P_k_*(0) = 1, when *k* = 0 and *P_k_*(0) = 0, otherwise.

### Simulating cluster formation dynamics

To understand amount of data needed to accurately estimate parameters of mathematical models describing cluster formation, we performed stochastic simulations of T cell clustering using Gillespie algorithm [17]. Simulations were run using `NestWhileList` function in Mathematica applied to the `OneStep` function (**Supplemental Figure S1**). In our models, change in the number of T cells in the cluster around the parasite is driven by the parameters λ_*k*_ (entry rate) and *μ_k_* (exit rate) with specific forms being different in different models of cluster formation. For example, in the random birth-death model, *λ_k_* = λ_0_ and *μ_k_* = *kμ*, and simulating cluster formation in this model showed stochastic but consistent increase in the number of T cells around individual parasites – assumed to be independent (**Supplemental Figure S2**A). As expected, the number of T cells found around individual parasites at a fixed time after simulation start followed a Poisson distribution [12], and the average cluster size increased over time as predicted by the model (**Supplemental Figure S2**B-C). This suggests that stochastic simulations are accurately describing the dynamics of cluster formation. Note that these simulations allow to track the number of T cells per parasite over time (longitudinal (LT) data) as well as to sample the number of T cells per parasite for different parasites at a specific time point (cross-sectional (CS) data).

We have simulated dynamics of cluster formation for 1,000 parasites with following parameters (unless noted otherwise): DDR model: λ_0_ = 0.2/h, λ_1_, = 0.2/h, and *μ* = 0.15/h; DIE model: λ= 0.8/h, *μ* = 0.7/h. These are typical parameter values found when fitting these models to the experimental data [12, 16]. While simulations provide a continuous change in the number of T cells for individual parasites over time, it is more typical experimentally to record the number of T cells present near the parasite at fixed time points. In our further analyses we sampled trajectories of T cell cluster size around individual parasites at different time points, most typical being 2, 4, and 8 hours after T cell transfer (and start of the clustering process).

### Statistics

To fit mathematical models to CS clustering data, we maximized the following likelihood function

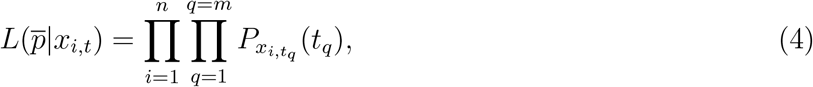

where *P_x_i,t_q___* the probability to observe a cluster of size *x_i,t_* for *i*^th^ parasite in the sample (*i* = 1,2… *n*) at time points *t_q_* as predicted by a given model and 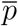 is the vector of model parameters (e.g., 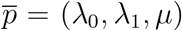 for the DDR model). We will call this method of fitting models to the data as CSL method since in these data we disregard the cluster size evolutions around individual parasites [12].

In contrast, to fit mathematical models to LT data, we maximize the probability that T cell cluster size will follow a particular path as observed in the data for individual parasites when clusters are measured for the same parasite at different times *t_q_* [16]. The likelihood of the model given the

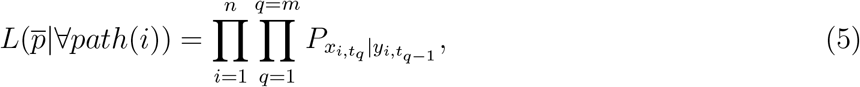

where *P_x_i,t_q___*|*y_i,t_q–1__* is the probability that for *i*^th^ parasite, *x* T cells were observed at time *t_q_* given that at previous time *t_q–1_ y* T cells were observed and 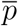 are the model parameters [16]. When solving the mathematical model numerically, transition probabilities are determined by solving ODEs from time *t*_*q*–1_ to *t_q_* assuming that *x* and *y* T cells, respectively, were present at the parasite site; *x* and *y* values are then taken from the experimental observations. We will call this the LTL method. We typically find model parameters by minimizing negative log-likelihood, 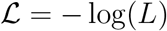.

Given that the DDR model was the best at describing clustering Plasmodium-specific T cells for most datasets [16], we calculated the accuracy and precision of the parameters of the DDR model found using the CSL and the LTL methods when we assumed different sampling times and number of parasites sampled per time point. Accuracy was defined as the difference between the true mean and the estimated mean, and the precision was defined as the coefficient of variation (CV) that is the square-root of the variance of the estimates over the estimated mean. To compare alternative models fitted to the CS data we used AIC and Akaike weights [18].

## Results

### Analytical solution of the DDR model explains why fitting the model to one time point data does not allow to estimate all model parameters

In most of our earlier analyses of the T cell clustering data we focused on predicting the steady state distribution of the cluster sizes and fitting these model predictions to data [12, 16, **Figure 1**]. The rationale for using a steady state approximation was that there was a statistically significant but small change in cluster sizes between 4 and 8 hours after the T cell transfer suggesting that most of cluster formation occurred early [16]. However, the steady state distributions of different models were defined by a combination of model parameters, thus, precluding the ability to estimate these parameters individually.

Fitting numerical solutions of the best fit DDR model to cross-sectional (CS) data of T cell clusters at 4 hours post T cell transfer revealed that the model is overdefined precluding convergence to a unique set of parameters [16]. To understand the reasons for this we used standard methodology [19–21] to find an analytical solution of the DDR model (see Supplemental Information for more detail):

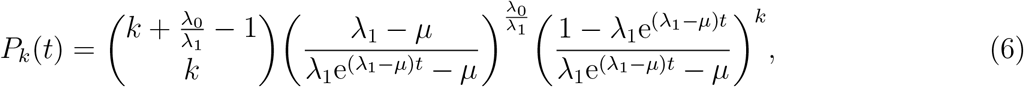

which has a form of a negative binomial distribution as a function of time *t*, of which *α* = λ_0_/λ_1_ is the parameter yielding the number of successes, and 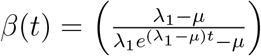 is the parameter yielding the probability of successes in a single trial of sampling *k* at a given time *t* (**Supplemental Figure S3**).

While the DDR model has 3 parameters, a closer inspection shows that its analytical solution (eqn. (6)) only depends on two parameter combinations, λ_0_/λ_1_ and λ_1_/*μ* (or equivalently, λ_0_/*μ* and λ_1_/*μ*). This provides the direct explanation why fitting the numerical solution of the DDR model to clustering data collected at one time point did not converge [16]. A formal analysis (see Supplemental information for more detail) proves this point further and suggests that measurement of T cell clusters at least two time points (at *t* > 0) is needed to estimate all model parameters, i.e. the model is identifiable [22].

While the alternative, density-independent exit (DIE) model did not fit the clustering data well we nevertheless found its analytical solution following previous calculations [23, and see Supplemental information for more detail]:

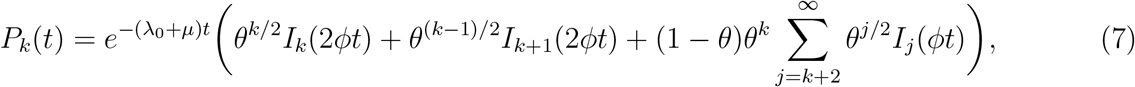

where *θ* = λ_0_/*μ*, 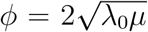 and *I_k_* is the modified Bessel function of the first kind. Interestingly, the analytic solution of the DIE model, in contrast, yields the number of functional parameters as same as the number of biological parameters, λ_0_ and *μ*. Therefore, measuring T cell clustering at one time point after T cell transfer is sufficient to estimate the two parameters of the DIE model (λ_0_ and *μ*), i.e., this model is also identifiable [22].

The result that the number of time points at which clustering of T cells is recorded influences ability to estimate all parameters of the DDR model is important. First, from one time point measurement all model parameters cannot be determined but the ratio of the parameters [16]. Second, when comparing alternative models, misspecifying the number of fitting parameters will bias AIC values by 2, and may in error reduce the Akaike weight of the DDR model in the list of alternative models. We should note, however, that while we did not know exact reasons we were aware that only 2 parameters of the DDR model can be estimated from the available CS data, and this was included in AIC calculations when comparing alternative models [16].

### Accuracy and precision in parameter estimates of the DDR model fitted to CS vs LT data

We performed stochastic simulations to more rigorously evaluate the impact of the following experimental details on accuracy and precision in estimating parameters of the DDR model (which was the best model found in our recent analysis [16]): 1) the number of measurement times (2 or more), 2) the number of parasites per time point (10-100), and 3) the method of data collection (CS vs. LT). All these details can be selected in experimental design but they have different costs and logistics, and knowing the minimal requirements may be extremely valuable to plan future (final) experiments.

In the simulations we generate the number of T cells found near individual parasites over time which then can be sampled to mimic our previous or future experimental design (**Supplemental Figures S1 and S2**). As it was expected from analytical results, sampling T cell cluster trajectories at one time point and fitting the DDR model to these data resulted in inadequate convergence and parameter estimates (results not shown). Fitting the DDR model to the simulated data sampled at least 2 time points revealed several interesting results.

First, even though stochastic simulations were performed with the same parameters of the DDR model, fitting the DDR model to the individually simulated datasets resulted in a wide range of estimated parameters with the average being closer to the true values when measurements were done at 3 time points (**Supplemental Figures S4 and S5**). Perhaps unsurprisingly, estimates were more variable when the models were fitted to CS data than when they were fitted to LT data, and this was true even for a larger number of parasites sampled (**Supplemental Figure S6**). Second, a more detailed analysis suggested that the number of parasites sampled per time point had the major impact on the difference between estimated and true parameter values (that we defined as accuracy) with the difference between estimates found for CS vs. LT data becoming relatively small at larger number of parasites sampled (**Figure 2**). Accuracy was higher when measurements were done at 3 time points, independently of the data type used (**Figure 2**). There was a positive bias in estimated model parameters at lower n and negative bias at higher n with its absolute value being smaller at a larger number of measurements per time point; LT data provided less biased estimates as compared to CS data (**Figure 2**).

**Figure 2:**
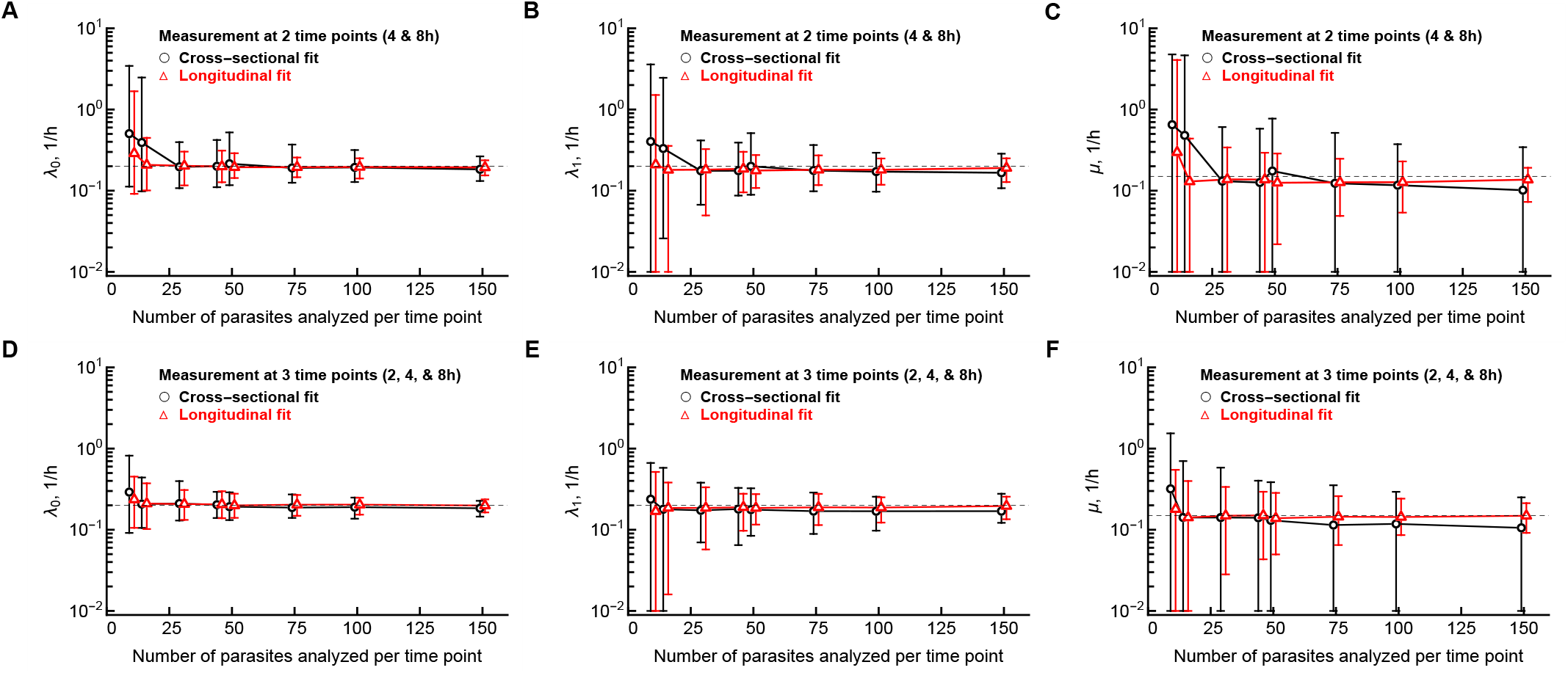
Accuracy of parameters of the DDR model estimated using cross-sectional (CSL) vs longitudinal (LTL) methods. We simulated the dynamics of cluster formation for 10^3^ parasites and sampled different numbers *n* of these simulated cluster size trajectories for 2 (4 and 8 hours; panels A-C) or 3 (2, 4, and 8 hours; panels D-F) time points after T cell transfer (see Materials and methods and **Supplemental Figure S4** for detail). We fitted the DDR model to the sampled CS or LT data using likelihood methods (eqn. (4) and eqn. (5), respectively) and estimated its parameters. Analysis was repeated 100 times for a given *n* and sampling time interval. Parameters of the DDR model used in simulations were λ_0_ = 0.2/h, λ_1_ = 0.2/h, and *μ* = 0.15/h and are denoted by thin dashed horizontal lines on individual panels. The error bars denote the 95% confidence intervals from 100 estimated values; note that due to plotting on log-scale the lower bound for the CI was fixed at 10^-2^.

Third, there was a large difference in how the number of parasites sampled influenced precision at which parameters were estimated; we defined precision as coefficient of variation (CV). Specifically, while precision in estimating recruitment rates became high (or error low) with increasing sample size, exit rate from the cluster could not be estimated well even when measurements were performed at 3 time points (**Figure 2** and **Supplemental Figure S6**). The larger confidence intervals were in part because many model fits of the CS data resulted in estimated exit rate from the cluster μ to be very close to 0 suggesting that such “simulated” data can be explained without T cell exit from the cluster (**Figure 2**C&F). This suggests that future studies may need not to focus on estimating the exit rate precisely but on defining the upper boundary of the exit rate that is consistent with the data. Importantly, the LT data were best suited to estimate the exit rate from the cluster due to relatively small errors (**Figure 2**C&F and **Supplemental Figure S6**C&F); however, obtaining relatively precise estimates may require tracking T cell clustering for 3 time points (~ 8 hours) for 100-150 parasites which may be prohibitively expensive or ethically impossible. Taken together, this analysis suggests that using cross-sectional data collection would require 3 time points and 50 parasites per time point for a relatively accurate and precise estimation of the parameters of the DDR model — with the caveat that the exit rate from the cluster may still have relatively low precision (**Supplemental Figure S6**). Collecting cluster formation details longitudinally (LT data) would be preferred because this would allow to more accurately estimate all model parameters including the exit rate *μ* with a smaller sample size; however, it remains to be determined if it is possible ethically (e.g., to keep mice under a microscope for 8 hours).

One important conclusion in our analysis so far is that LT data are typically more informative than CS data (although more difficult to collect). However, our comparisons between the two types of data were not fully compatible because the simulated data were generated for different parasites. Therefore, we performed another set of simulations in which we simulated T cell cluster formation using the DDR model but this time we selected the same parasites (*n* = 32 or *n* = 50) and sampled them at 3 time points (2, 4, and 8 hours after T cell transfer) but then treated the sampled data as CS (and thus, as independent parasites/samples) or as LT and fitted the DDR model to these data using appropriate likelihood functions (eqns. (4)–(5)).

As found in our other results we observed a large variability in estimates of the DDR model between individual runs further emphasizing stochastic nature of modelled cluster formation (compare **Figure 3** and **Supplemental Figures S4 and S5**). Surprisingly, there was a statistically significant but overall poor correlation between values of the parameters found by fitting the DDR model to the data from the same parasites using CSL vs. LTL methods; for example, there were examples when LTL method provided a relatively large estimate of the T cell exit rate from the cluster while CSL method provided near zero estimate (**Figure 3**C). As expected, sampling more parasites per time point reduced variability in the parameter estimates for recruitment rates and less so for the exit rate. Interestingly, we found that the LTL method on average provided higher and more accurate estimates of the parameters, and that the CSL method typically underestimated the true rates. These results further emphasize superiority of the LTL method in estimating parameters of the DDR model but also illustrate that parameter estimates may differ substantially between individual “datasets” suggesting a need to perform several independent experiments.

**Figure 3:**
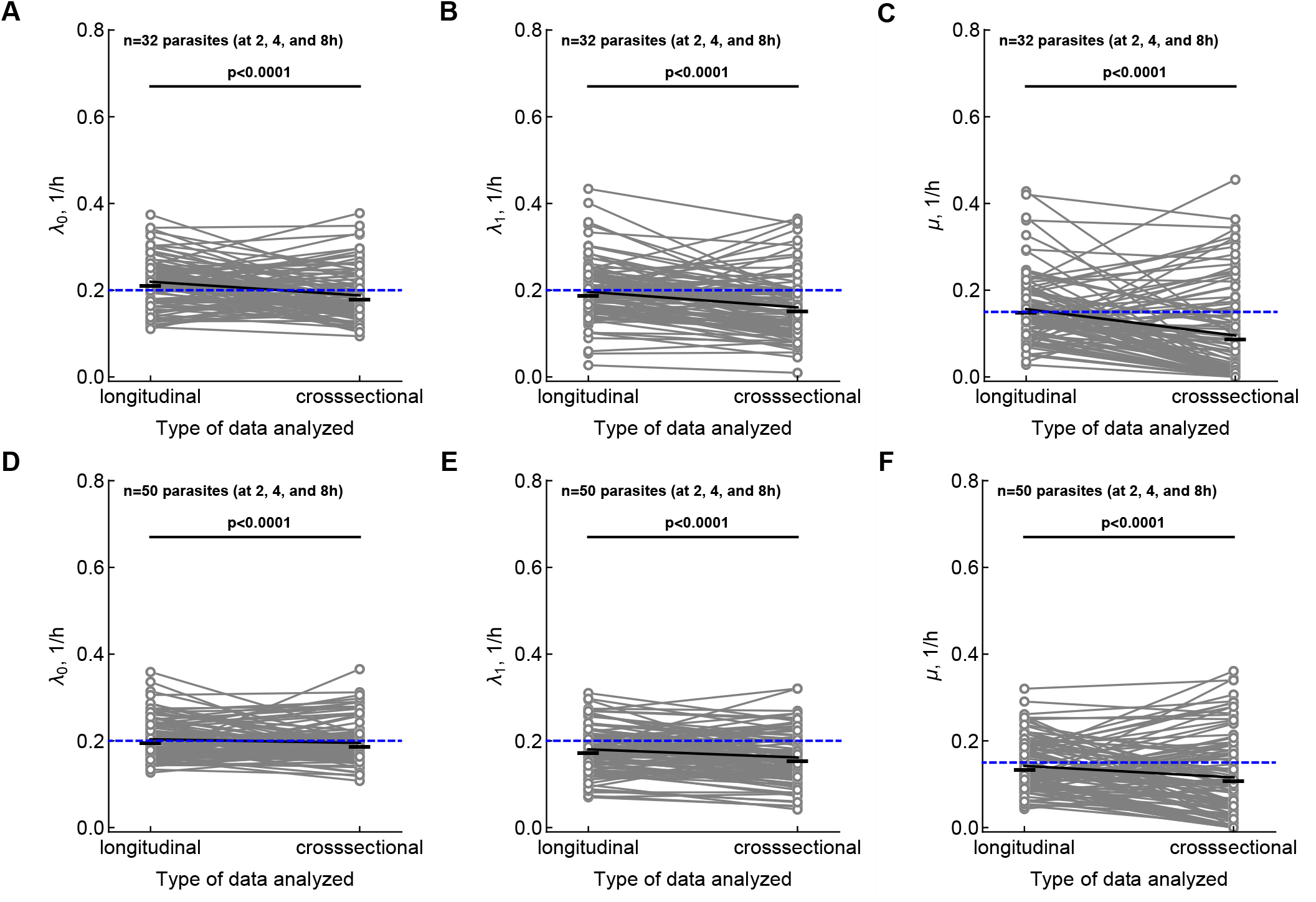
Variability in parameter estimates of the DDR model found CSL vs. LTL estimation methods for the same sampled trajectories. We simulated the dynamics of cluster formation using the DDR model as previously (**Figure 2**), sampled *n* = 32 (panels A-C) or *n* = 50 (panels D-F) parasites at 3 time points after T cell transfer (2, 4, and 8 hours). We then fitted the DDR model to the simulated data assuming that the data are CS or LT. Because the model was fitted to effective the same T cell cluster trajectory, parameter estimates are paired (as indicated on the panels by gray lines). Black solid lines denote the average parameter estimate and horizontal dashed blue lines denote the parameter values assumed in simulations (λ_0_ = 0.2/h, λ_1_ = 0.2/h, and *μ* = 0.15/h). Analysis was repeated 100 times. Medians were compared using signed rank test with p-values from the test indicated on individual panels.

### Moderate data size is required to robustly discriminate between DDR and DIE models

In our previous analyses we have shown that the DDR model could fit the data on clustering of Plasmodium-specific CD8 T cells in murine livers much better than any other alternative models tested [16]. However, the DDR model still failed to explain some experimental observations, for example, formation of very large clusters within 4 hours since T cell transfer suggesting that other mechanisms of cluster formation may be at play (for example, different parasites may display different levels of “attractiveness” for T cells). An alternative DIE model could describe some of the clustering data although in most cases with lower quality. We wondered if there is a critical size of the data that one needs to collect to robustly discriminate between the DDR and DIE models. To address this question we performed stochastic simulations.

Specifically, we simulated T cell cluster formation in accord with either DDR or DIE models, sampled different number of parasites/trajectories per time point (*n* = 32, *n* = 50, or *n* = 80) for 3 time points (2, 4, and 8 hours) post T cell transfer assuming “independent” parasites (i.e., CS data) and fitted the DDR or DIE models to these data using CSL method. Then we compared the quality of the model fits to these simulated data using Akaike weights *w* (**Figure 4**). Perhaps not surprisingly, the model that was used for stochastic simulations typically fitted the data better than the alternative model as is indicated by a *w* > 0.5 for such fits. However, in many instances the alternative model provided a better fit; for example, DIE model fitted the simulated data, generated using the DDR model, better than the DDR model in 12/100 cases for *n* = 32 parasites sampled (**Figure 4**A). Increasing the number of parasites sampled naturally resulted in smaller number of better fits by an alternative model, and at *n* = 50 parasites sampled per time point (for 3 time points), the frequency of model misclassifcation was lower than 95% (**Figure 4**B&E). Interestingly, while simulating T cell clustering with the DDR model and sampling *n* = 80 parasites resulted in very large weights for the DDR model fits, the opposite was not observed (**Figure 4**C&F). In fact, the DDR model fitted many simulated data, generated using the DIE model, with somewhat reasonable quality (with w = 0.2 – 08). This suggests that the DDR model is sufficiently flexible to fit the data that have a different underlying mechanism. Indeed, the DDR model has 3 parameters while the DIE model has only 2 to be estimated from the data. This result suggests that even if the DDR model fits the experimental data with best quality, other, independent data are needed to provide additional support for the model [1].

**Figure 4:**
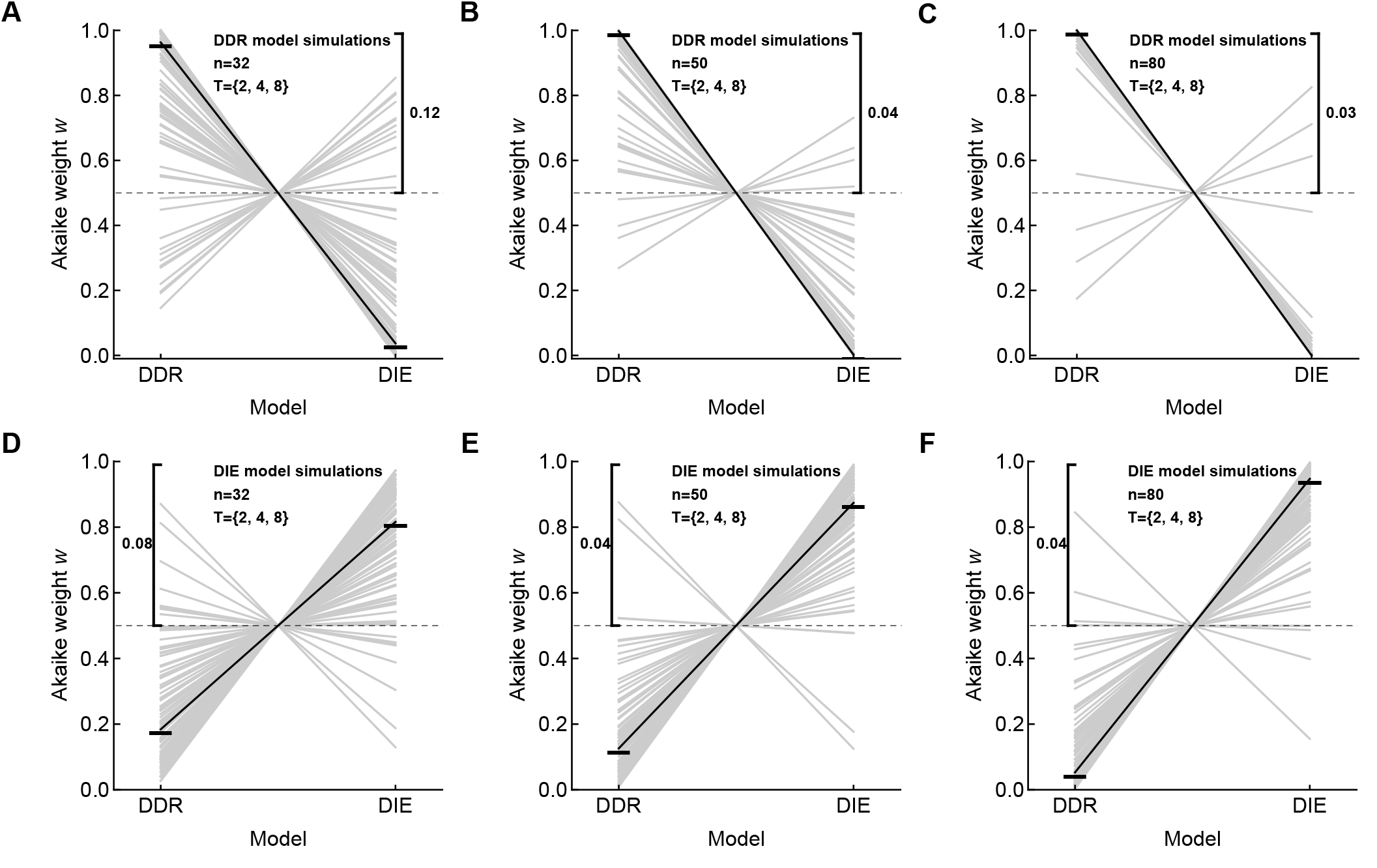
A moderate number of sampled parasites is needed to discriminate between alternative models when using CSL method. We simulated cluster formation in accord with the DDR model (panels A-C) or DIE model (panels D-F), sampled 32 (A&D), 50 (B&E), or 80 (C&F) parasites at 2, 4, and 8 hours post T cell transfer. We then fitted the DDR or DIE models (**Figure 1**) to these simulated data assuming that these are CS data (eqn. (4)) and compared the quality of the two alternative model fits using Akaike weights *w*. Black lines denote average *w* per model, and the frequency of model fits with *w* > 0.5 when the alternative model was fitted to the data is indicated on individual panels. We performed 100 sets of simulations with parameters for the DDR model being λ_0_ = 0.2/h, λ_1_ = 0.2/h, and *μ* = 0.15/h and for the DIE model being λ = 0.8/h, and *μ* = 0.7/h.

### General framework to guide experimental design with mathematical modeling

Based on our specific example of CD8 T cell clustering around Plasmodium liver stages we propose a general framework of how mathematical modeling can be used to help design future experiments (**Figure 5**). Specifically, this framework should be used to perform power analyses, i.e., to to provide estimates of the size and type of the data that are required to accurately estimate parameters of mathematical models and/or to discriminate between alternative models.

**Figure 5:**
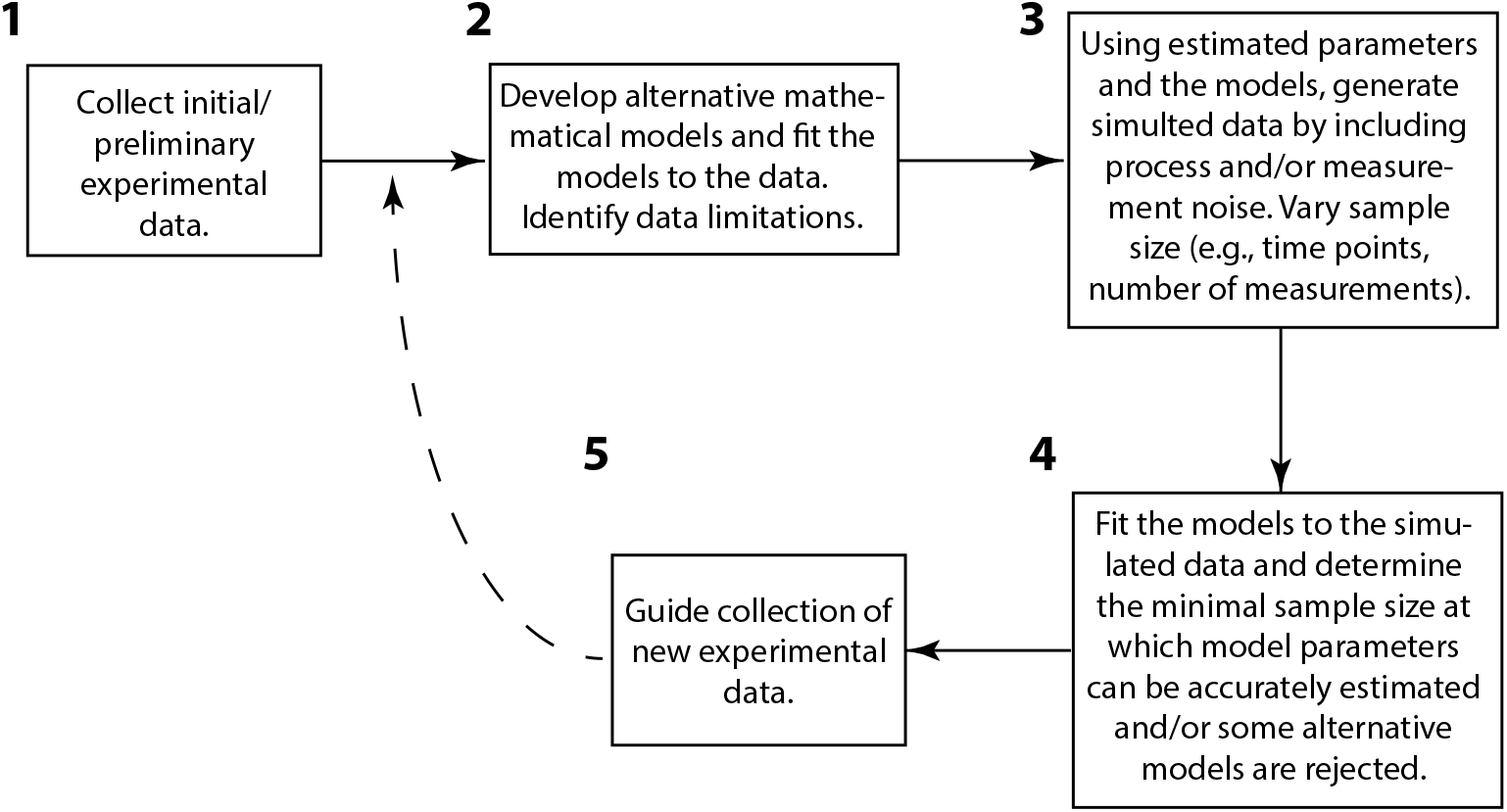
General framework to use mathematical modeling to guide experimental design. Understanding a given biological phenomenon typically begins with collection of preliminary data (step 1). Given the data and other details about the biological system in question one can develop alternative mathematical models and fit the models to data (step 2). Here the limitations of the data may be identified, e.g., if the data are of insufficient size to estimate all model parameters and/or to discriminate between alternative models. Using quantitative insights generate so far (e.g., estimates of model parameters), one can simulate the biological system in question and generate simulation/synthetic data assuming each model is true by allowing randomness in the process (process noise) and/or in measurements (measurement noise, step 3). In step 4, the models are re-fitted to the simulated data, allowing one to determine the threshold size and the type of the data that discriminates between the alternative models. In step 5, one makes recommendations of the size and the type of data to the experimenters, given their constraints on cost and time-to-collection functions. The cycle may be repeated (by going to step 2) especially if new experiments revealed better understanding and new alternative models for the biological system in question.

The framework begins with collection of experimental data about the system in question (step 1 in **Figure 5**). The type of such preliminary data may vary in type and size but typically need to be quantitative in nature, to allow for fitting the models to the data. In our experience, the data collection often happens before mathematical models of the underlying biological processes are thought of. For example, many of the data on clustering of Plasmodium-specific CD8 T cells in the liver (i.e., number of T cells per every imaged parasite) had been collected before we developed alternative mathematical models of the clustering process [12]. The type of the data and biological expertise of the experimentalists should help with development of appropriate mathematical models for the biological system in question [1]. In step 2, the models are fitted to the data and potential limitations of the data are identified. For example, the amount of the data may not be sufficient to estimate all the model parameters. This was the case for the DDR model fitted to the T cell clustering data collected at one time point. Furthermore, the data may be not sufficient to discriminate between alternatives (e.g., [5]); this was not the case in the T cell clustering data that did allow to provide stronger statistical support for the DDR model [16]. Nevertheless, fitting models to data may still provide estimates for some model parameters and allow to put bounds on other parameters. These estimates can be then used in step 3 to generate synthetic/simulation data. There may be different ways of generating such simulation data. One approach can be to run Monte Carlo simulations to allow for randomness in the biological process (i.e., process noise) as we did for T cell clustering (**Supplemental Figures S1 and S2**). An alternative approach is to simulate model dynamics deterministically and add some degree of measurement noise to the model predictions. Magnitude of the measurement noise could be estimated from the data or assumed as a part of the power analysis. A key feature of this step 3 is to generate simulated data of different sizes by varying the number of samples per dataset, which components of the biological system can be sampled, etc. In step 4, the models are fitted to the simulated data and deviation of the estimated parameters from the parameters assumed in simulations (e.g., accuracy of the estimates) can be evaluated as the function of the generated or assumed noise in the data and the amount of data collected. The same simulated data can be used to determine the size of the data allowing to discriminate between alternative models. The two goals – estimating model parameters and discriminating between alternative models — may require different sizes or types of data. Finally, in step 5 results of the analysis lead to recommendations of the type of data that needs to be collected to estimate model parameters and/or to discriminate between alternative models. This step should also consider different costs of experiments including time and consumables needed (**Figure 5**).

## Discussion

Developing alternative mathematical models for underlying biological processes and disseminating between such alternatives with experimental data is an important component of rigorous science [1]. However, the size of the data that may be needed to discriminate between alternatives as well as to accurately and precisely estimate parameters of the best fit models remains rarely estimated in immunology (but is typically required in NIH grant applications). By using a specific example of CD8 T cell clustering around Plasmodium-infected hepatocytes we illustrated how stochastic simulations can be used to perform a power analysis and to determine the impact of sampling (number of parasites sampled, time points at which sampling is performed, and the way of data collection) influences estimates of the model parameters and allows to discriminate between alternative models. This approach requires that some preliminary data are collected to guide development of initial mathematical models and estimation of their parameters. This situation is similar to clinical trials in which initial trials (typically phase 2) allow to estimate a potential effect size of a given treatment; these data are then used to perform power analyses for the size of larger, phase 3 clinical trials to detect the expected effect size.

Using an analytical solution of the DDR model we showed that at least 2 time points are required to estimate its parameters while one point is sufficient for the DIE model parameter estimation. Although the DDR model is mean- and median-unbiased for increasing parasite numbers in a sample, there is, however, a systematic positive error in the estimates at small parasite numbers with fewer time points in the data (**Figure 2**). Analysis of the longitudinal data (LTL method) yields more accurate and reliable estimates of the DDR model parameters as compared with the cross-sectional data (CSL method) at small sample sizes (**Supplemental Figure S6**). As a thumb rule we propose using a minimum of *n* = 50 parasites if one uses the cross-sectional data collection method, and *n* = 30 if one uses the longitudinal data collection method allowing both yielding comparable confidence intervals. Measurements of cluster size should be taken at least at three time points of data provide reasonable confidence intervals for model parameters (**Supplemental Figure S6**). The confidence intervals of the exit rate *μ* in the DDR model get bloated in either case, but LTL method can keep them markedly tighter than the CSL method. A parasite sample size of more than *n* = 50 does not seem to improve the distinguishability between the DDR vs the DIE models in cross-sectional analyses for the analyzed parameters of the models (**Figure 4**). The bias in the mean estimates of the DDR model at small parasite sample sizes (**Figure 2**) and with fewer number of time points may arise from the statistical structural feature in the DDR model solution, which is a time-dependent negative binomial distribution, skewed heavily cross-sectionally at early time-points (see eqn. 6) and Supplemental information for detail). Our study also suggested the possibility of miss-identifying the degrees of freedom in the DDR model by assuming that its three parameters can be estimated from one time point CS data. Such miss-specification will increase the AIC by 2 that may result in selection of the incorrect alternative model as the best fit model.

As far as we are aware our results are relatively unique examples of how power analysis of a complex biological process may be performed. And yet, our study has several limitations. First, while we performed extensive analyses on the number of parasites one need to sample to reliably estimate model parameters, we did not exhaustively explored the parameter space for the DDR or DIE models and the time points. In part, this is driven by constraits of our specific experimental system for which we have already estimated model parameters (e.g., [12, 16]). We do not expect that in our future experiments best fit model parameters will be drastically different. Similarly, we focused on specific times at which T cell clustering has been (or can be) measured driven again in part by our experimental collaborators who suggested these specific time points. However, researchers interested in extending our work to their own biological systems involving clustering of T cells should repeat similar analyses with parameters that are more reasonable for their circumstances. Such parameter values should be driven by the preliminary experimental data available for analysis. While we have provided analytical solutions of the DDR and DIE models that will speed up fitting these models to experimental data, the solutions in their current form cannot be used in analysis of the LT data. Solutions that are generalized for a cluster size *k* at time *t* = 0 will need to be developed. Furthermore, these analytical solutions assume that model parameters are constant in time. Our recent work suggested that the rates of T cell entry into the cluster and T cell exit from the cluster may be declining with time [16]. Whether analytical solutions of these models with time-dependent parameters exist remains to be determined. While the fitting the DDR model to the LT data yields more robust parameter estimates, this procedure is computationally expensive because it requires to solve a system of ODEs for every path of the T cell cluster size observed in the data [16]. Increasing the number of time points will also increase this cost substantially. Finally, our previous experimental design in which mice were initially infected with Plasmodium sporozoites and then received infusion of Plasmodium-specific CD8 T cells are not fully representing prophylactic vaccination. Our more recent experiments involve transfer of T cells first that is followed by infection with Plasmodium sporozoites [24]. Given that sporozoites spend only minutes in the blood [25, 26], our power analyses should be directly applicable to this new, more biologically-relevant experimental design.

Our work suggests an important use of mathematical modeling in immunology – by using data from preliminary experiments to perform power analyses to direct and justify the size of future (final) experiments (**Figure 5**). This is clearly is an important part of rigorous science. Predictions on the sample size and time points at which data should be collected can be then tested in experiments that hopefully will improve our understanding of how vaccine-induced CD8 T cells locate and ultimately eliminate Plasmodium liver stages in mice.

## Abbreviations

DDR: density-dependent recruitment
DIE: density-independent exit
LT: longitudinal
CS: cross-sectional
LTL: longitudinal likelihood
CSL: cross-sectional likelihood
NBD: negative binomial distribution.

## Data sources

No data were used in the paper.

## Codes

Simulations have been performed either in Matlab (version 2018b) or Mathematical (version 11.2 or 12.3). Simulation codes will be made available upon request.

## Author’s contributions

Mathematical analyses and analytical proofs were done by HR. Simulations of cluster formations were done primarily by VVG. HR wrote the first draft of the paper with all authors contributing to the final draft.

## Acknowledgements

This work was supported by the NIH (R01GM118553 and R01AI158963) awards to VVG.

## Supplemental Information

Evaluating impact of experimental design on accuracy of parameter estimation and model selection efficiency: T cell clustering as a case study

### Analytical solution of the density-dependent recruitment (DDR) model

Let the immigration term λ_0_ be the probability at which the cluster size increases linearly from one state to an immediately higher state, independent of the cluster size of the present state, λ_1_ be the per capita probability of transition of cluster size from one state to a higher state, and *μ* be the per capita probability of cluster size decreases from one state to the immediate lower state. Thus, we the DDR model is given by the following equations

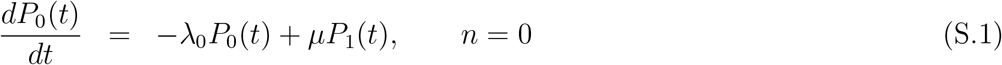

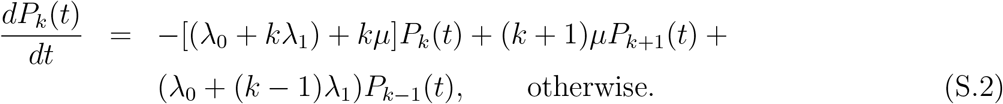

Assuming that *μ* > A_1_, the steady state solution for this model was calculated previously [12]. For this linear model, the probability generating function (PGF) technique can be applied for writing down the appropriate partial differential equations for the PGF [21]. An explicit solution to eqns. (S.1)–(S.2) can be obtained by solving them via moment generating function:

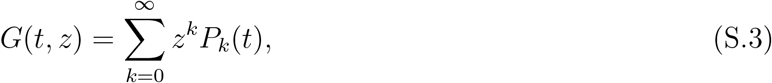

where *z* is a dummy variable. Multiplying both sides of eqns. (S.1)–(S.2) by *z^k^* and summing over *k*, and substituting the following partial derivatives we find

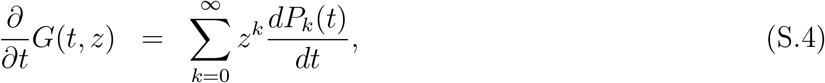

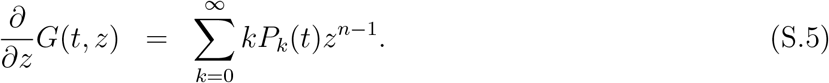

Re-indexing the summations from *k* = 1 to *k* = ∞ into *k* = 0 to *n* = ∞ without loss of generality we get the following Lagrangian type partial differential equation:

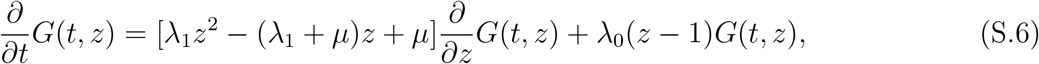

which is identical to eqn. 7) in chapter 2 of Goel & Richter-Dyn [21]. We solve eqn. S.6) using the following auxiliary equation:

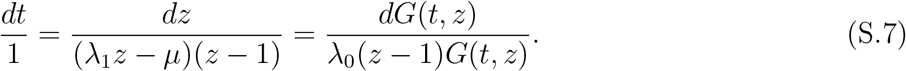

We take the first and second terms of eqn. S.7),

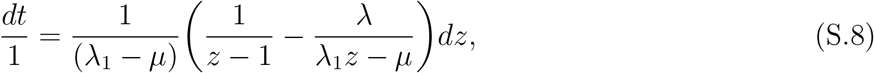

which yields 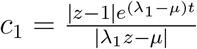 where *c*_1_ is a constant. From second and third expressions of eqn. S.7) we find

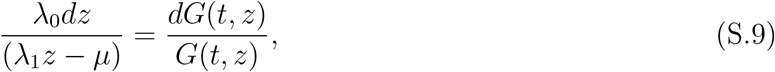

which yields 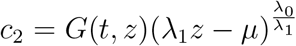 where *c*_2_ is a constant. Thus, from eqn. S.9) we can write the general solution of eqn. S.6) as

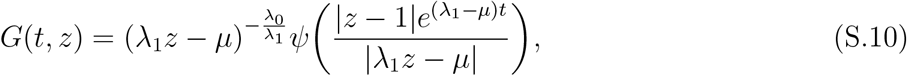

where *ψ* is yet an unspecified function. By substituting initial condition *G*(*z*, 0) = 1 in eqn. S.10), and taking *P*_0_(0) = 1 and *P*_>0_(*t* = 0) = 0 in eqn. S.10) we get

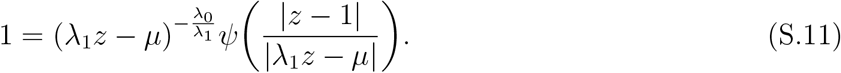

By taking 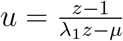 we get 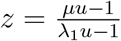 and substituting it in eqn. S.11), we get

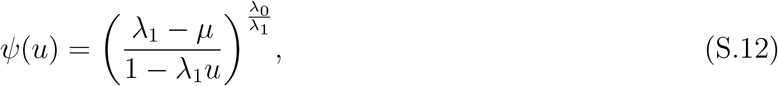

and substituting 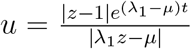 in eqn. S.12), eqn. S.10) can be written as the following moment generating function.

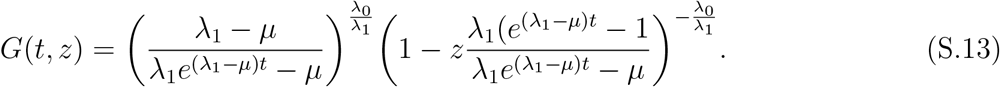

Let’s choose 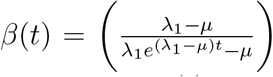. By choosing *γ*(*t*) = 1 – *β*(*t*) we get 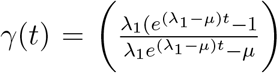.

Note that for *t* = 0, we get *β*(*t*) = 1 and *γ*(*t*) = 0. And for *t* → ∞, we get *β* = 0 and *γ* = 1. By taking 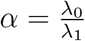 and substituting *β*(*t*) and *γ*(*t*) in eqn. S.13) we find

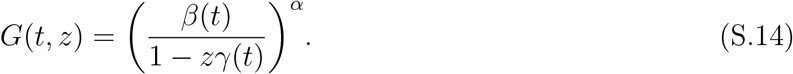

Note that (1 – *zγ*(*t*))^-*α*^ can be expanded as a series 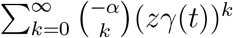 or equivalently as

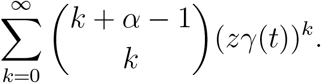

Thus, we can write eqn. S.14) as

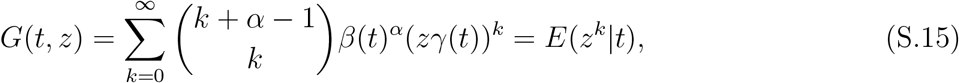

where *k* is the cluster size. This can be derived similarly, had we used the form *z* = *e^θ^* also, where 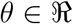. From eqn. S.15), we can easily show that the derivatives of *G*(*t, z*) with respect to *z* give as *G*′(*t, z*) = *β*(*t*)^*α*^ *αγ*(*t*)(1 – *γ*(*t*)*z*)^-*α*-1^ and *G*′′(*t, z*) = *β*(*t*)^*α*^*α*(*α* + 1)*γ*(*t*)^2^(1 – *zγ*(*t*))^-*α*-2^and so on.

Thus, by general theory, solving *k*^th^ derivative of the moment generating function *G*(*t, z*) at *z* = 0 yields *P_k_*(*t*) as follows:

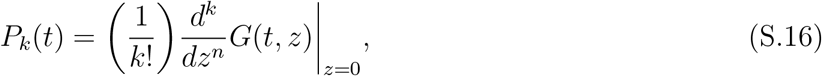

resulting in the following time-explicit solution for *P_k_*(*t*), which satisfies the initial conditions *P*_0_(0) = 1 and *P*_>0_(0) = 0:

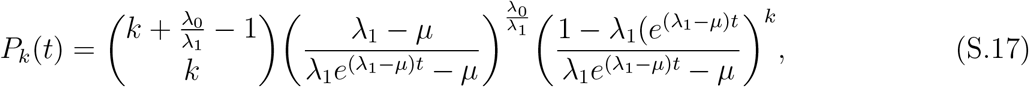

which is identical to the solution found in Renshaw [20]. The expected value of *k* at time *t* can be found as 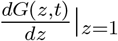. It yields

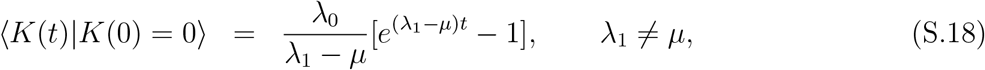

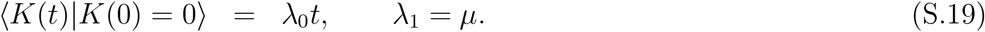

And similarly, any *n*^th^ factorial moments can be derived by

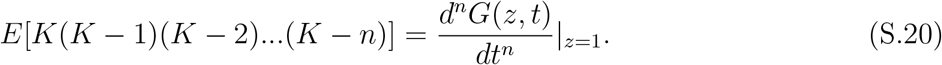

This solution has a stable probability distribution of cluster sizes *k* for *t* → ∞ from eqn. S.15). For λ_1_ < *μ*

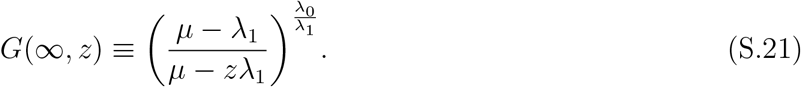

Thus, from eqn. S.16)

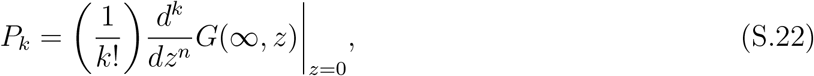

which are time-independent solutions. In summary, birth-death-immigration process yields time-dependent negative binomial distribution as a solution, with two parameters *NB*(*k*, *α*, *β*(*t*)), (equivalent to standard notation *NB*(*k*, *n*, *p*)), where as the numerical solution remains with three parameters as per the original ODE for of the DDR model. The continuous approximation to this is a Gamma distribution (**Supplemental Figure S3**).

### Analytical solution of the density-independent exit (DIE) model

The density-independent exit (DIE) model is given by the following differential equations:

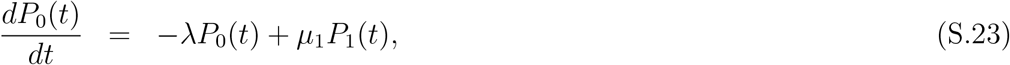

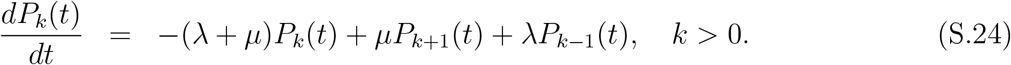

The steady state solution of the model for *μ* > λ was found in our previous study [12]. The equation of moment generating function *G*(*t,z*) is then given by

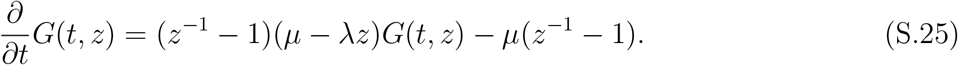

We Laplace-transform the above equation to find

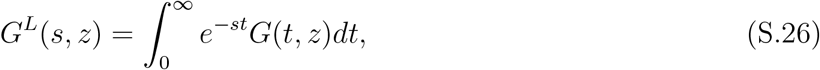

the property 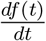 transforming to 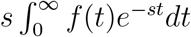, and Laplace transform of *G*_0_(*t*) as

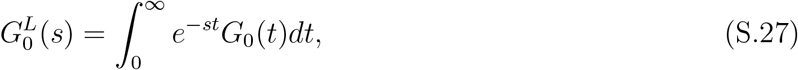

obtaining

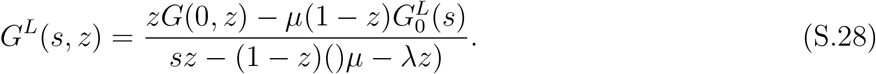

By definition

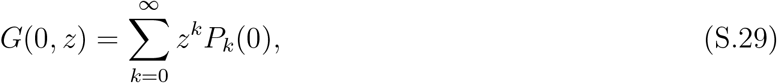

and by taking initial conditions *P_k_*(0) = 1 for *k* = 0 and *P_k_*(0) = 0 for *k* > 0 we find

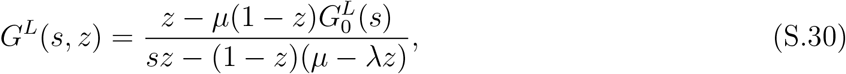

The 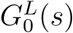 is determined through denominator roots of the above equation, and taking the doubly Laplace inverse transformation we find the model solution:

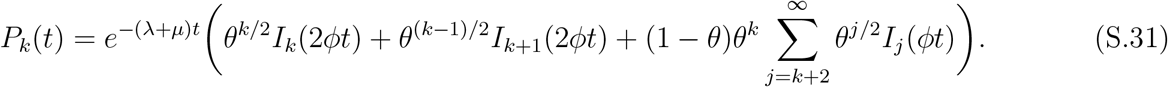

### Measurement at least two time points are needed to estimate parameters of the DDR model

From eqn. S.17), the solution of the DDR model can be written as the negative binomial distribution (NBD) with the binomial coefficients in the definition of the pdf being replaced by an equivalent expression as follows:

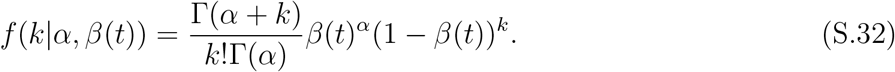

For the CS data, the likelihood function is given simply as

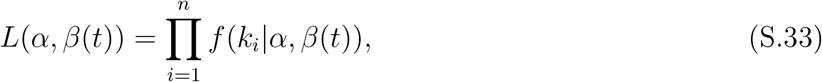

with the negative log-likelihood function given by

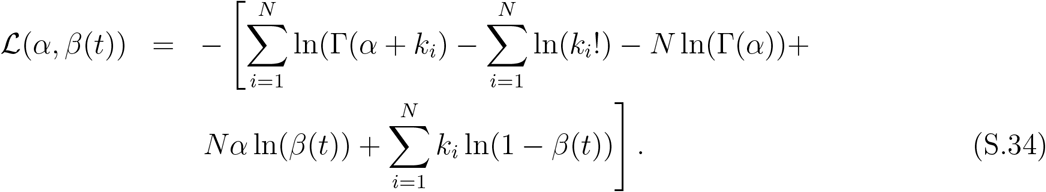

Setting the partial derivatives of the function with respect *α* and *β*(*t*) to zeros yields the maximum log-likelihood estimates of the parameters *α* and *β*(*t*):

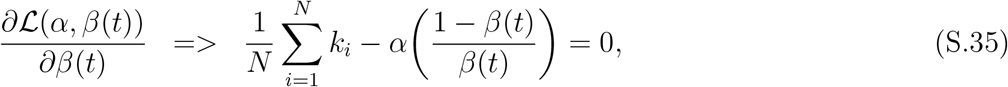

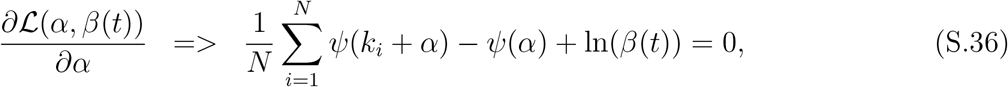

where *ψ* is a digamma function. From eqn. S.35) we get

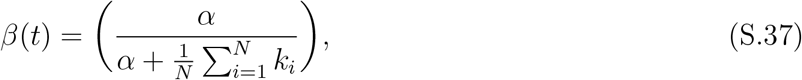

where 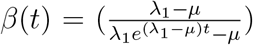. By substituting eqn. S.37) in eqn. S.36) we get an exclusive function of α as

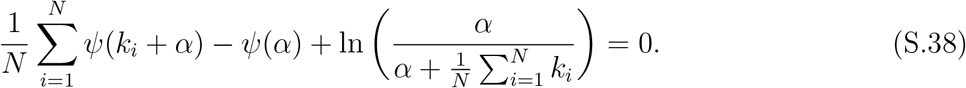

Equation (S.38) has a unique solution for *α*, but not closed form, yet, can be solved using an iterative technique such as Newton’s method. Hence, we take the solution as the constant *c* as below:

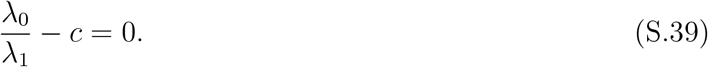

Furthermore, from the previous result by substituting 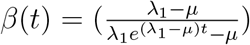 in eqn. S.35) we can show that

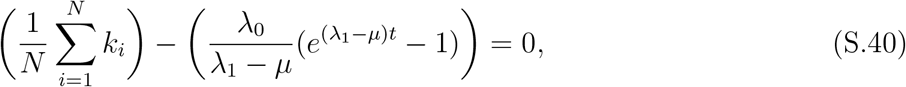

which can be written as

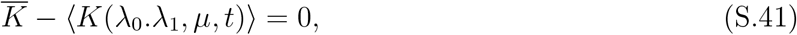

where 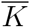 is an arithmetic mean and 〈*K*(*t*)〉 is the mean drift of the process at time *t*. That is, for example, if t is a constant for all data, as in the example below, and *n* = 30, it can be easily seen that the average of the data *k_i_* equals 〉*k*(*t* = 20)〉 in eqn. S.18). This can be computed numerically.

#### Measurement of cluster sizes at one time point

From eqn. S.39) we get a unique value (estimate) for *α* = *c*, and thus substituting it in eqn. S.37), we get a unique fixed value (estimate) for *β*(*t* = *t*_1_), resulting from the maximum likelihood of the estimation of 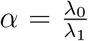, and 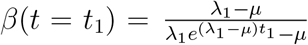. Equation (S.40) can also be written as

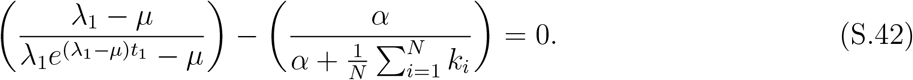

Equations (S.39) and (S.40) (and eqns. (S.41)–(S.42)) cannot be solved for unique values of λ_0_, λ_1_, and *μ*. These parameters are interrelated, conditioned by the unique fixed values of *α* and *β*(*t* = *t*_1_) at the maximum likelihood estimates. Therefore, for the data that contain only measurement of cluster sizes at one time point, all parameters of the DDR model cannot be uniquely determined. Furthermore, one can easily show that the mean-drift function 〈*K*(*t*)〉 can take any trajectory, conditioned by the intercepts at (0,0) and *k* = 〈*k*(*t* = 20)〉 at time *t* = *t*_1_. Taken together, these results prove that data sampled at a single time point are insufficient to estimate parameters λ_0_, λ_1_ and *μ* uniquely; instead, parameters *α* and *β*(*t*) (or ratios λ_0_/*μ* and λ_1_/*μ*) can be accurately estimated.

#### Measuring the cluster size at two different time points

Assume that we have *n*_1_ number of data *k*^1^ from one time point *t*_1_, and *n*_2_ number of data *k*^2^ from another time point *t*_1_, s.t., *t*_1_ < *t*_2_. In this case, the likelihood function can be written as

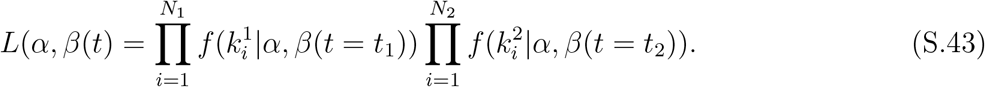

Thus, the negative log-likelihood function is

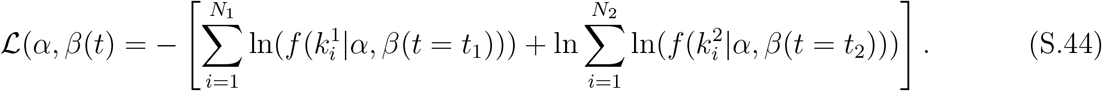

Thus, following the methods from above the following holds true:

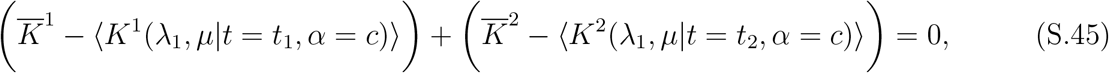

where 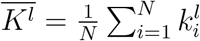 and 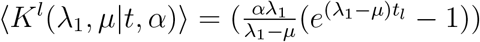, where *l* = 1, 2 indicate data from time points *t* = *t*_1_ and *t* = *t*_2_ respectively, and *c* is the unique estimate of 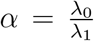. Here, *α* can be solved for unique value iteratively yielding a constant. Thus, we take the solution for *α* and compute *β*(*t*). Thus, it is mathematically equivalent to the case where eqn. S.42) has only two parameters to be solved for; λ_1_ (or instead λ_0_) and *μ*. Thus, it can be easily seen that there is a unique solution exists for λ_1_, and *μ* in eqn. S.45) taking 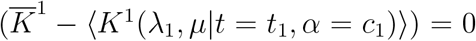 and 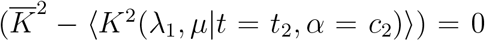 if (〈*K*^1^〉) < (〈*K*^2^〉) for *t*_1_ < *t*_2_. Assume that there are other estimates of λ_1_ and *μ* that satisfy eqn. S.45), we can write

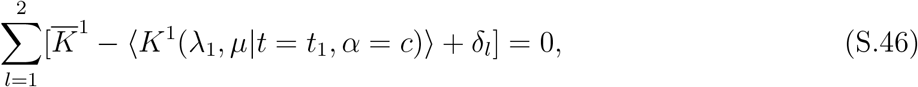

where *δ_l_* are small deviations from the fit of mean data from the mean estimations. Equation (S.46) yields unique solution to λ_1_ and *μ*, estimated at 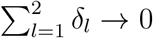. Furthermore, note that in the case of having data from more than 2 time points, given that at least one set of measurements are from a time point of clustering evolution, we get

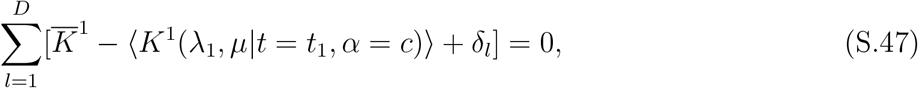

where *D*(> 2) are the the time points of the data. Here also, note that *α* is time-independent, thus unique. Thus, it allows eqn. S.47) to estimate λ_1_ and *μ* similarly to the case before, yielding unique solutions.

**Supplemental Figure S1:**
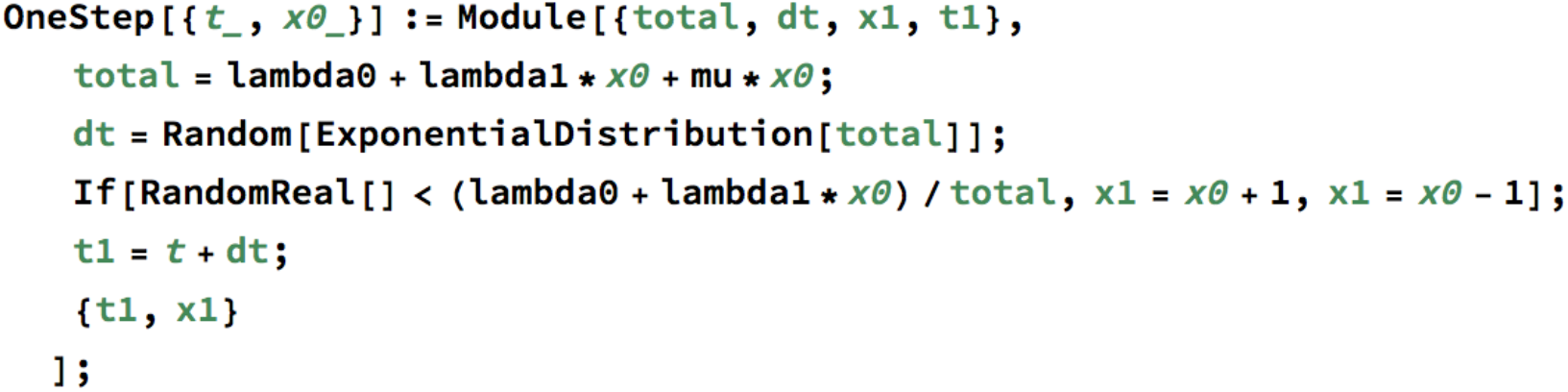
The basic Mathematica code to simulate Gillespie stochastic dynamics of cluster formation. The provided example is for the random birth-death model (when λ_1_ = 0) or for density-dependent recruitment (DDR) model (when λ_1_ > 0, see eqns. (1)–(3) in Main text). In simulations *dt* is the random time when a reaction occurs, and the second random number determines whether the cluster grows in size or is reduced in size by one at frequency proportional to the rate of these two processes. Another function OneStepDIE was developed to simulate the cluster formation dynamics for density-independent exit (DIE) model.

**Supplemental Figure S2:**
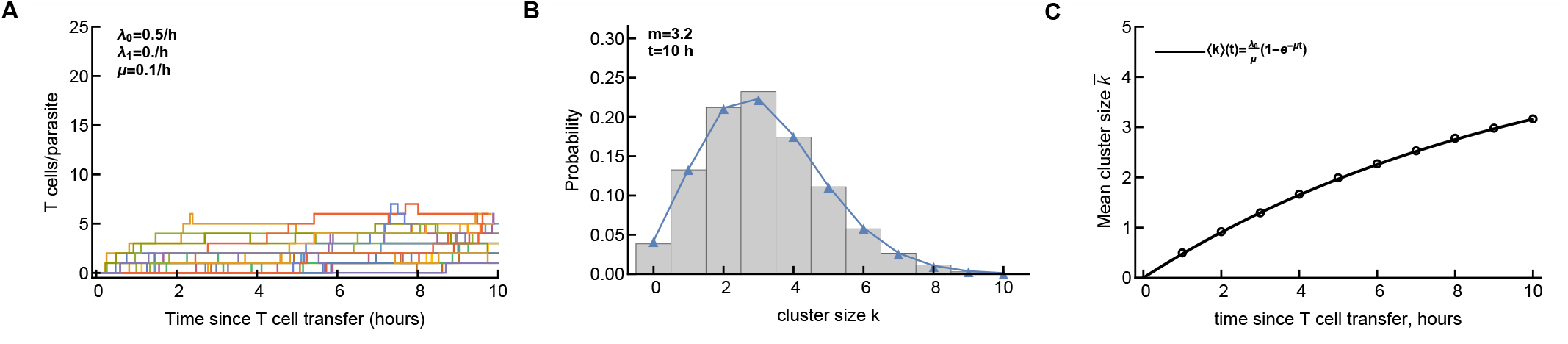
An example of stochastic simulations of the CD8+ T cell cluster formation. Here we assume random entry-random exit model with λ_0_ = 0.5/h, λ_1_ = 0, and *μ* = 0.1/h. Panel A shows 20 individual runs, panel B shows the distribution of cluster sizes at 10 hours after start for 5 × 10^3^ simulations, and panel C shows the change in the average cluster size in simulations (points) and as predicted by the random birth-death model.

**Supplemental Figure S3:**
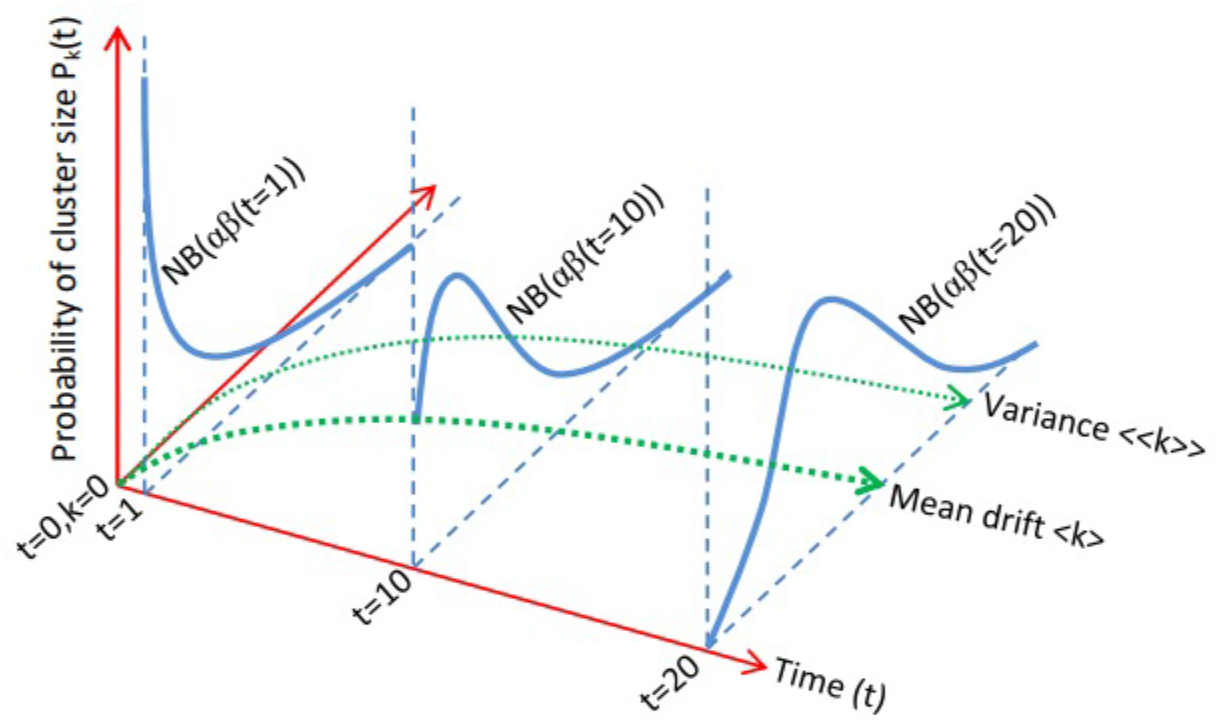
Dynamics of the DDR model. Time-evolution of probabilities of the random birth-death-immigration process (eqn. S.17)) that stats from the delta function (at *t* = 0) through approximating exponential, gamma, and ultimately normal distributions over time, arising from time-dependent negative binomial distribution, *NB*(*k*, *α*, *β*(*t*)), with cluster size *k* and parameters *α* and *β*(*t*) as function of time.

**Supplemental Figure S4:**
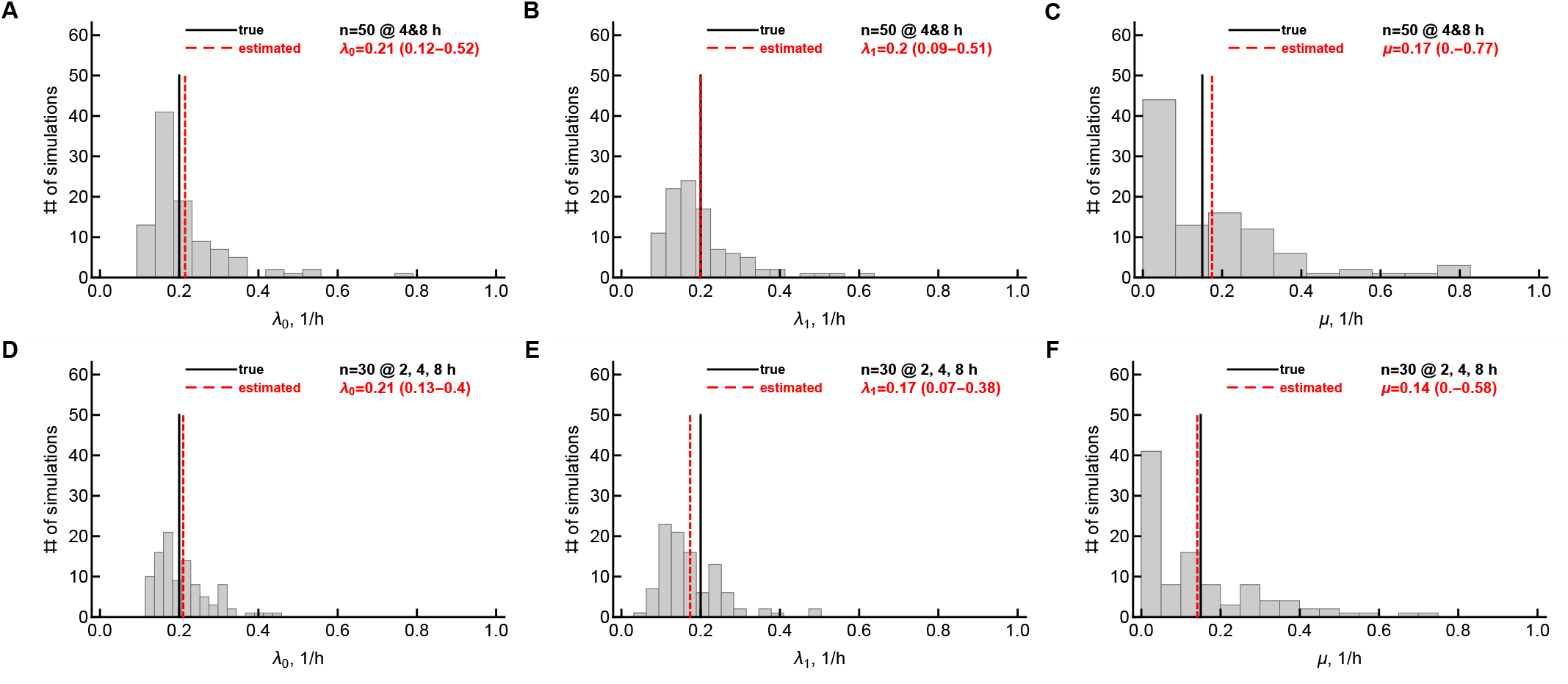
The parameters of the DDR model have a wide distribution when the model is fitted to CS data. We simulated the dynamics of T cell cluster formation assuming that it is driven by the DDR model with parameters λ_0_ = 0.2/h, λ_1_ = 0.2/h, *μ* = 0.15/h (**Supplemental Figures S1 and S2**) and sampled the simulated data at 2 time points (4 and 8 hours after the start with *n* = 50 parasite/trajectories sampled per time point, panels A-C) or 3 time points (2, 4, and 8 hours after the start with *n* = 30 parasites/trajectories sampled per time point, panels D-F). Different number of parasites sampled per time point was to limit the overall sampling size to about 100 parasites per experiment; sampling was done by randomly choosing trajectories from simulated data per time point, thus, representing CS data. We fitted the DDR model to each of the datasets and its parameters (λ_0_, λ_1_, and *μ*) were estimated using CSL method (eqn. 4)). The procedure was repeated 100 times and resulting distribution of estimated parameters of the DDR model are shown on individual panels. Vertical lines denote the true parameter value (solid back lines) and estimated value (dashed red lines) for individual parameters, and 95% confidence intervals are shown on individual panels in parentheses.

**Supplemental Figure S5:**
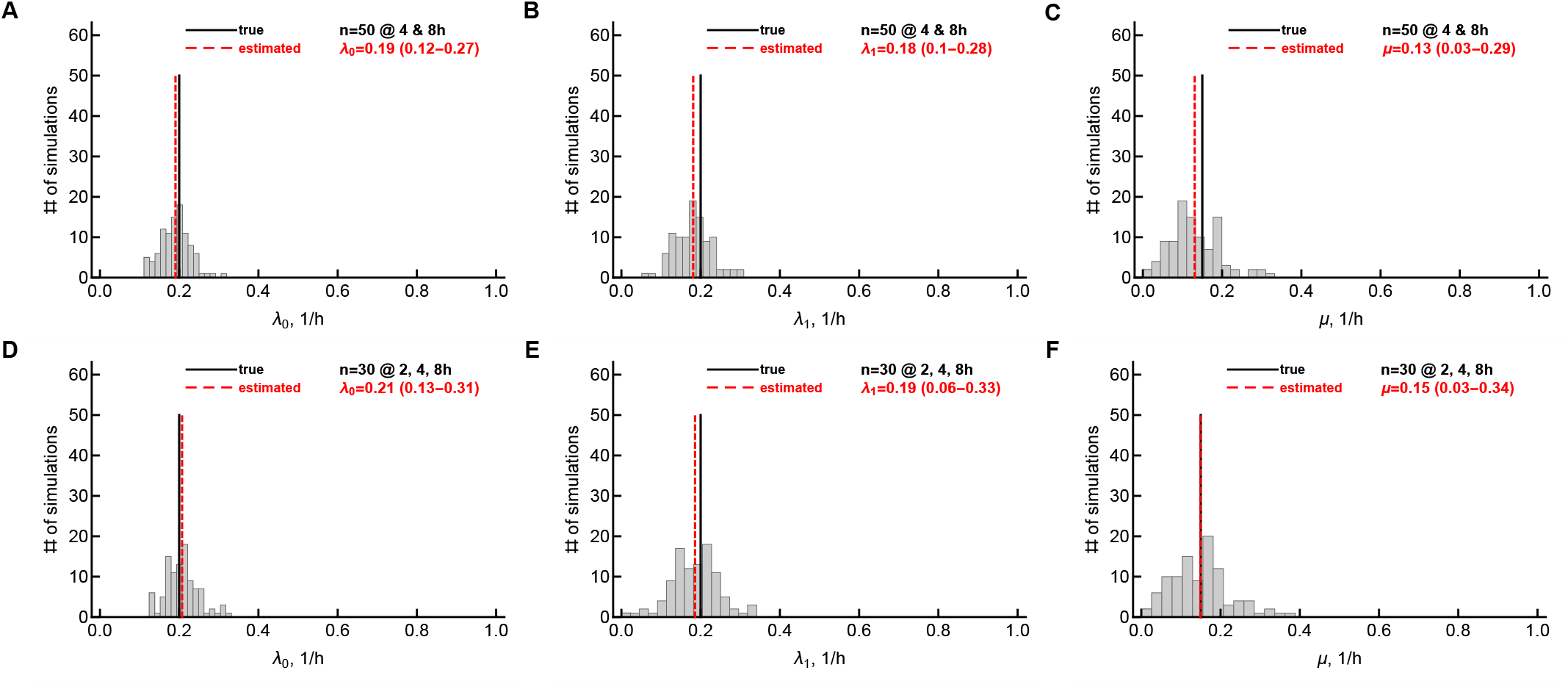
The parameters of the DDR model have a relatively narrow distribution when the model is fitted to LT data. We performed the same simulations as in **Supplemental Figure S4** but instead sampled T cell cluster sizes for individual parasites/trajectories over time, i.e., longitudinally, representing, thus LT data. We fitted the DDR model to each of the datasets and its parameters (λ_0_, λ_1_, and *μ*) were estimated using LTL method (eqn. 5)). The rest of notations are the same in as **Supplemental Figure S4**.

**Supplemental Figure S6:**
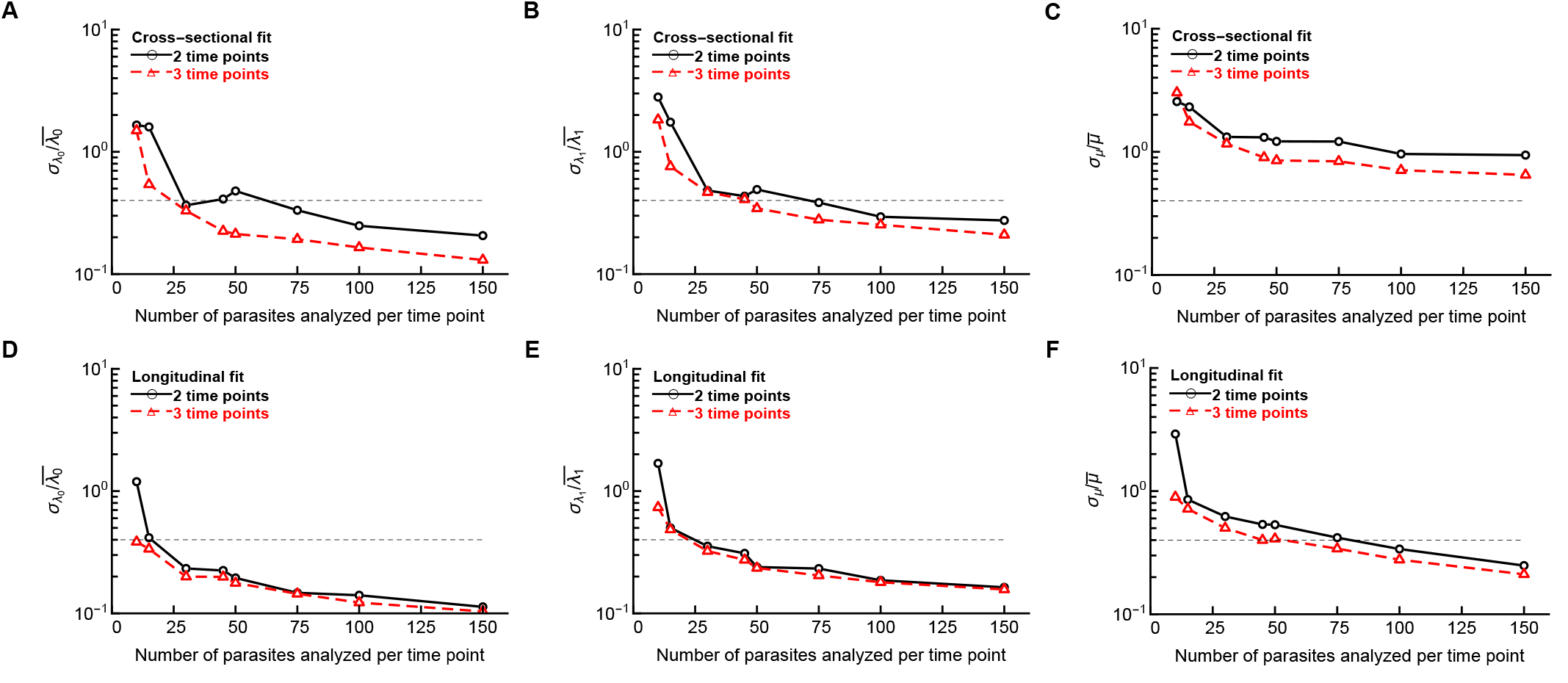
A smaller number of parasites needs to be tracked longitudinally than cross-sectionally to provide accurate estimates of the DDR model. We simulated the T cell clustering dynamics as described in **Supplemental Figure S4** and sampled from the simulations the different number of parasites surveyed per time point assuming either CS (panels A-C) or LT (panels D-F) types of data. Two time points samples (solid black lines with circles) were done at 4 and 8 hours post T cell transfer and three time points samples (dashed red lines with triangles) were done at 2, 4, and 8 hours post T cell transfer. The DDR model was fitted to these data, and the coefficient of variations (CV = standard deviation/mean) of the estimated parameters are shown on individual panels. Horizontal dashed lines denote CV = 0.4.

